# A long interval between priming and boosting SARS-CoV-2 mRNA vaccine doses enhances B cell responses with limited impact on T cell immunity

**DOI:** 10.1101/2022.08.03.502672

**Authors:** Alexandre Nicolas, Gérémy Sannier, Mathieu Dubé, Manon Nayrac, Mark M. Painter, Rishi R. Goel, Mélanie Laporte, Halima Medjahed, Justine C. Williams, Nathalie Brassard, Julia Niessl, Laurie Gokool, Chantal Morrisseau, Pascale Arlotto, Cécile Tremblay, Valérie Martel-Laferrière, Andrés Finzi, Allison R. Greenplate, E. John Wherry, Daniel E. Kaufmann

## Abstract

Spacing the first two doses of SARS-CoV-2 mRNA vaccines beyond 3-4 weeks raised initial concerns about vaccine efficacy. While studies have since shown that long-interval regimens induce robust antibody responses, their impact on B and T cell immunity is poorly known. Here, we compare in SARS-CoV-2 naïve donors B and T cell responses to two mRNA vaccine doses administered 3-4 versus 16 weeks apart. After boost, the longer interval results in higher magnitude and a more mature phenotype of RBD-specific B cells. While the two geographically distinct cohorts present quantitative and qualitative differences in T cell responses at baseline and after priming, the second dose led to convergent features with overall similar magnitude, phenotype and function of CD4^+^ and CD8^+^ T cell responses at post-boost memory timepoints. Therefore, compared to standard regimens, a 16-week interval has a favorable impact on the B cell compartment but minimally affects T cell immunity.

## INTRODUCTION

The standard SARS-CoV-2 mRNA vaccine regimens recommend an interval of 21-day (Pfizer-BioNTech BNT162b2) or 28-day (Moderna mRNA-1273) between vaccine doses. However, the optimal interval has not been determined in controlled trials. In the context of vaccine scarcity and given the significant protection already conferred by the first dose in non-high-risk populations (Baden et al., 2021; Polack et al., 2020; Skowronski and De Serres, 2021), some public health agencies implemented schedules with longer intervals to rapidly extend population coverage (Paltiel et al., 2021; Tuite et al., 2021). While such strategies generated concerns given uncertain immunogenicity, a longer period of partial vulnerability to infection and a hypothetical risk of escape mutant selection, epidemiological evidence supports this approach as a valid alternative in lower-risk populations (Carazo et al., 2021; Skowronski et al., 2021) in which robust T cell and antibody responses are observed after a single dose (Tauzin et al., 2021b). Recent reports suggest that an extended interval between priming and boost procured enhanced humoral responses (Grunau et al., 2021; Hall et al., 2022; Payne et al., 2021b; Tauzin et al., 2021a).

As protective antibodies are associated with vaccine efficacy (Earle et al., 2021; Gilbert et al., 2021), there is a need to better understand the generation and maintenance of B cell memory responses elicited by different vaccine modalities. As CD4^+^ T cell help provided by T follicular helper (Tfh) is critical for the expansion, affinity maturation and memory development of B cells (Crotty, 2019), it is also important to determine whether dosing interval affects CD4^+^ and CD8^+^ T cell vaccine responses. Demonstrating a direct protective role of SARS-CoV-2-specific CD4^+^ and CD8^+^ T cells independent of humoral immunity has been more challenging, but a number of studies support the notion that these lymphocyte subsets may contribute to recovery from COVID-19: Th1 cells, which foster development of CD8^+^ T cell memory (Laidlaw et al., 2016), and Th17 are important for mucosal immunity (Nayrac et al., 2022). However, T cell subsets show important heterogeneity and plasticity, better fitting with spectra of phenotypes and functions than fully distinct populations (O’Shea and Paul, 2010). Previous studies by our group (Nayrac et al., 2022) and others (Painter et al., 2021a; Rodda et al., 2022) have demonstrated temporal associations between vaccine-induced CD4^+^ T cell responses and development both B cell and CD8 T cell immunity that present partially different patterns depending on the interval between doses.

However, in contrast to the important progress made in the understanding of the kinetics of B and T cell responses in short interval mRNA vaccine schedules (Goel et al., 2021; Painter et al., 2021b; Rodda *et al*., 2022; Zollner et al., 2021), how a long interval between the first two vaccine doses affects B and T cell immunity compared to standard dosing regimens remains poorly known due to the paucity of studies performing side-by side comparisons with the same cellular immunity assays (Flaxman et al., 2021; Hall et al., 2022; Payne et al., 2021a).

Here, we apply standardized high-parameter flow cytometry assays to longitudinally compare the quantitative and qualitative features of vaccine-induced Spike-specific B cells, CD4^+^ T cells and CD8^+^ T cells in SARS-CoV-2 naïve participants enrolled in two cohorts: participants who received the two mRNA vaccine doses administered 16 weeks apart, defined as a long interval regimen; and participants who received the two doses 3-4 weeks apart, defined as a short interval regimen.

## RESULTS

### Study participants

We evaluated immune responses in two independent cohorts of health care workers (HCW) that received two doses of mRNA vaccines (Figure 1A). The two cohorts differed by the time interval between the priming and the boosting inoculations, which defined the long interval (LI) cohort (16-week spacing, n=26; Montreal cohort) and the short interval (SI) cohort (3-4 week spacing, n=12; Philadelphia cohort). Blood samples were examined at 4 time points: at baseline (B) before vaccination; 3 weeks after the first dose (D1); 1-3 weeks after the second dose (D2), and 10 to 16 weeks after the second dose (M2). Clinical characteristics are shown in Table 1. The median age of the participants in the short interval cohort was 15-year-old significantly younger (Mann-Whitney p = 0.019). Both cohorts significantly differed in the interval between prime and boost, and in the time of sampling D2 (3 weeks post second dose for LI, 1 week for SI) and M2 (16 weeks post second dose for LI, 10 for SI). No other statistical differences were noted.

**Table 1.** Clinical characteristics of the study participants^†^.

|  | Long Interval (LI)* | Short Interval (SI)* |
| --- | --- | --- |
| Prime-boost interval | 16 weeks apart | 3 weeks apart |
| Variable | (n=26) | (n=12) |
| Vaccine regimen |  |  |
| Pfizer BNT162b2 vaccine (2 doses) | n=25 | n=11 |
| Heterologous vaccine strategy (Moderna mRNA-1273 and Pfizer BNT162b2) | n=1 | n=0 |
| Moderna mRNA-1273 vaccine (2 doses) | N=0 | n=1 |
| Age (years) | 51 (41-56) | 38 (22-63) |
| Sex |  |  |
| Male | 11 (42%) | 4 (33%) |
| Female | 15 (58%) | 8 (66%) |
| Vaccine dose spacing (days) |  |  |
| Days between doses 1 and 2 | 111 (109-112) | 21 (20-28 ) |
| Visits for immunological profiling (days) |  |  |
| B, days before first dose | 1 (0-5) | 0 (-1-1) |
| D1, days after first dose | 21 (19-26) | 21 (20-28) |
| D1, days before second dose | 90 (85-92) | 0 (-1-0) |
| D2, days after first dose | 133 (130-139) | 29 (27-38) |
| D2, days after second dose | 21 (20-27) | 7 (7-12) |
| M2, days after first dose | 224 (222-228) | 94 (86-115) |
| M2, days after second dose | 112 (110-119) | 70 (65-94) |

<sup>†</sup> Values displayed are medians, with IQR: interquartile range in parentheses for continuous variables, or percentages for categorical variables.
\* The Long-interval (LI) and Short-interval (SI) cohorts were compared by the following statistical tests: for continuous variables, Mann-Whitney test, for categorical variables, Fisher's test.
Values highlighted in light grey are statistically different between the LI and SI cohorts.

**Figure 1.**
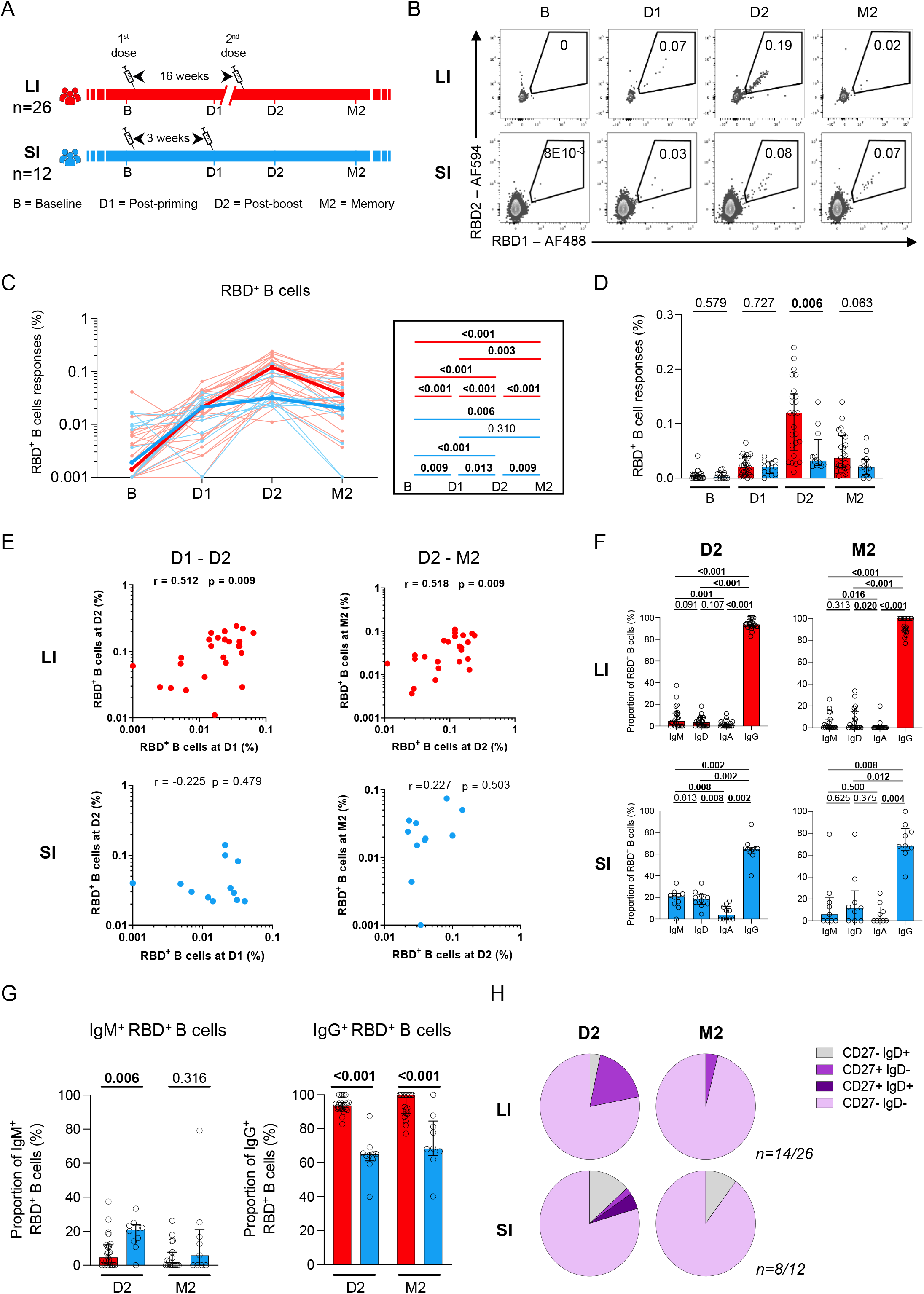
A 16-week delayed boost enhances the magnitude and maturation of B cell responses. **(A**) Schematic representation of study design. Blood samples were analyzed at 4 time points in the long (red) interval (LI) and short (blue) interval (SI) cohorts: baseline (B); 3 weeks after priming (D1), 1-3 weeks after boost (D2) and 10-16 weeks after boost (M2). **(B)** Representative examples of RBD-specific B cell responses. **(CD)** Kinetics of RBD-specific B cell responses in LI (red) vs SI (blue) cohorts. **(C)** The bold line represents cohort’s median value. Right panel: Wilcoxon tests. **(D)** Inter-cohort comparisons. Bars represent medians ± interquartile ranges. Mann-Whitney tests are shown. **(E)** Scatter plots showing temporal RBD^+^ B cell correlations in the LI and SI cohorts. r: correlation coefficient. Significant correlations by Spearman tests (p<0.05) are shown in bold. **(F)** Frequencies of IgD, IgM, IgA and IgG-positive cells within RBD-specific B cells within each cohort, paired comparisons with Wilcoxon tests. (**G)** Proportion of IgM^+^ and IgG^+^ cells among RBD^+^ B cell cells, with Mann-Whitney tests for comparison between the LI and SI cohorts. (**H)** Proportion of IgD^+/-^ and CD27^+/-^ populations in RBD-specific B cells. In **H**, only D2 and M2 provided enough events for analysis. In **C-E** n=26 for long-interval (LI), n=12 short-interval (SI). In **F-H** n=14 for long-interval (LI), n=8 short-interval (SI).

### A 16-week delayed boost enhances the magnitude and maturation of B cell responses

To evaluate SARS-CoV-2-specific B cells, we focused on the Receptor Binding Domain (RBD) of Spike to minimize inclusion of B cells cross-reactive to endemic coronaviruses (Hicks et al., 2021; Klumpp-Thomas et al., 2021). Co-detection of two fluorescently labeled recombinant RBD probes greatly enhances specificity (Figure 1B and (Anand et al., 2021) flow cytometry panel, Table S1; gating strategy, Figure S1A). We examined the magnitude of RBD-specific B cells (defined as RBD1^+^RBD2^+^CD19^+^CD20^+^) in the two cohorts (Figure 1CD). Most participants showed no signal at baseline, and clear RBD-specific B cell responses after priming that were very similar between the LI and SI cohorts at the D1 timepoint, as expected. In contrast, the second dose elicited brisk recall responses at D2 in the LI cohort whereas a plateau was observed in the SI cohort (Figure 1C), leading to statistically significant differences between LI and SI donors at this timepoint. The antigen-specific B cell responses subsequently declined at M2 in both cohorts, and while the relative decline was less pronounced in SI than LI participants, although a strong trend for higher memory responses in LI vaccinees persisted at M2 (Figure 1D). In contrast to short-interval participants, where no temporal association could be found between post-prime RBD^+^ B cell responses and post-boost RBD^+^ B cells, a strong and statistically significant positive correlation was observed in the long-interval cohort (Figure 1E). Likewise, RBD^+^ B cell responses at D2 were associated with stronger memory responses in the long-interval cohort (Figure 1E).

To determine whether the interval between vaccine doses qualitatively impacted antigen-specific B cell development, we measured IgM, IgD, IgA and IgG expression on RBD-specific B cells (Figure S1B). RBD-specific B cells in LI donors were almost entirely IgG^+^ at the D2 and M2 time points, contrasting with statistically significant lower IgG^+^ fractions in SI participants (Figure 1FG, Figure S1CD). Unswitched IgM^+^ and IgD^+^ RBD-specific B cells could be detected after boost in the SI cohort at both the D2 and M2 memory time points, with a statistically significantly higher fraction of IgM^+^ RBD^+^ B cell responses at D2 compared to LI donors (Figure 1FG).

To assess RBD-specific B cell differentiation, we next quantified IgD and CD27 co-expression (Figure S1E). CD27 is predominantly expressed on memory B cells (Tangye et al., 1998), and IgD on unswitched B cells (Moore et al., 1981). An atypical double-negative (DN) IgD^-^ CD27^-^ was dominant at both the D2 and M2 time-points in both cohorts (Figure 1H, Figure S1F). In the LI cohort, a class-switched memory IgD^-^CD27^+^ RBD-specific B cells was present at D2 and contracted at M2. This subset was negligible at both time points in the SI cohort. Only rare naïve-like IgD^+^CD27^-^ were observed in LI participants, while more IgD^+^CD27^-^ RBD-specific B cells were identified in the SI cohort at D2 and M2, (Figure 1H).

These data show that compared to the standard short interval regimen, the second vaccine dose given after a long 16-week interval elicits more robust and mature RBD^+^ B cell responses. A short 3-week interval results in a lower plateau of RBD^+^ B cell responses, associated with delayed maturation and weaker association between early post-boost and memory responses.

### The initial two-dose vaccination series elicits Spike-specific CD4^+^ T cell responses of similar magnitude irrespective of dosing interval

CD4^+^ T cells help play a critical role in development of B cell and CD8^+^ T cell immunity. We therefore measured Spike-specific T cell responses at the four time points in the two cohorts (Figure 2 and S2). As in our previous work (Tauzin *et al*., 2021b), we used both a TCR-dependent activation induced marker (AIM) assay that broadly identifies antigen-specific T cells and intracellular cytokine staining (ICS) to perform functional profiling (flow cytometry panels: Tables S2-3).

**Figure 2.**
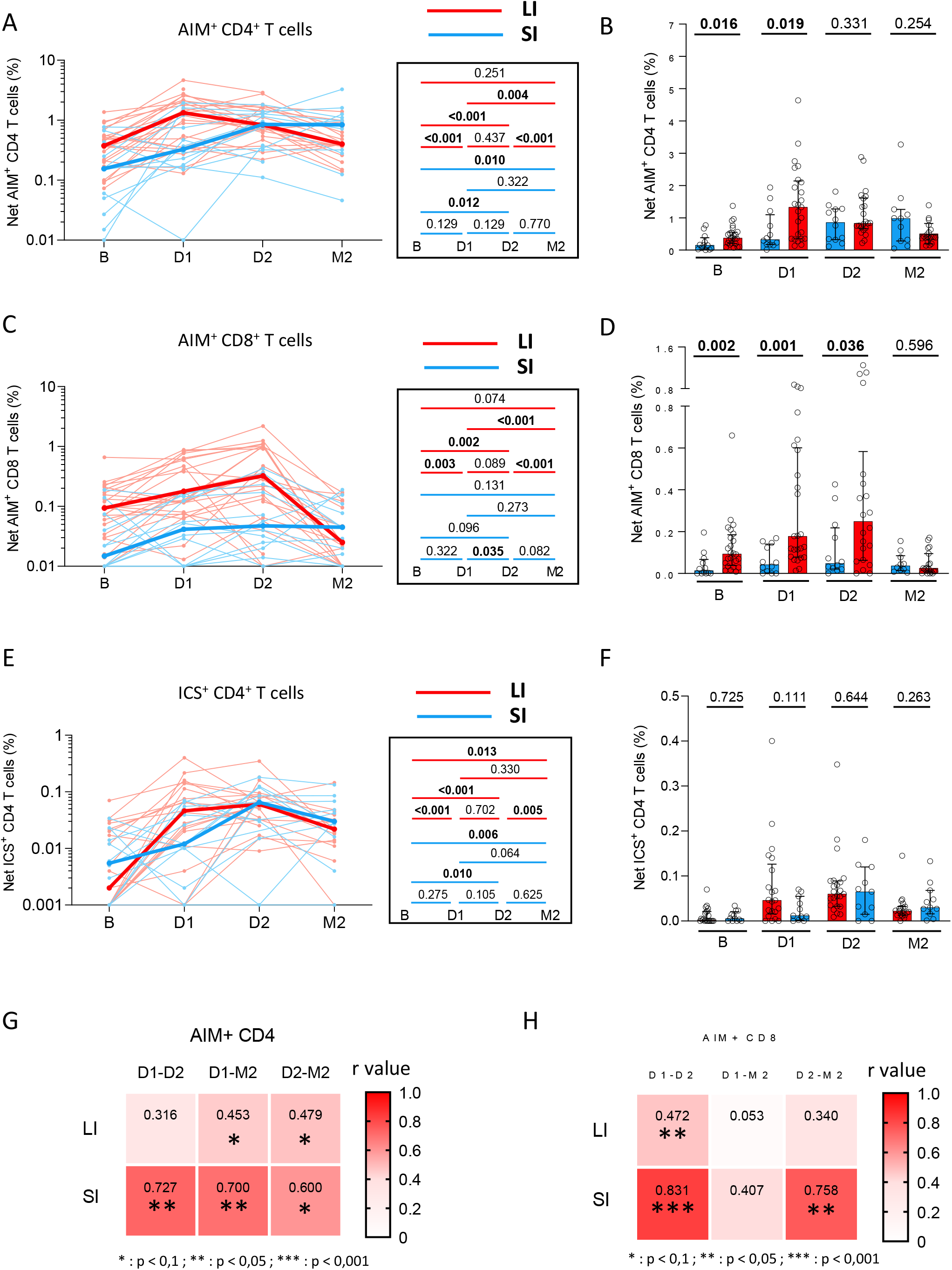
The initial two-dose vaccination series elicits Spike-specific CD4^+^ T cell responses of similar magnitude irrespective of dosing interval. SARS-CoV-2 Spike-specific CD4^+^ and CD8^+^ T cells in long (red) and short (blue) receiving two vaccine doses. **(AB)** Longitudinal (**A**) and inter-cohort (**B**) analyses of net Spike-specific AIM^+^CD4^+^ T cell responses. **(CD)** Longitudinal (**C**) and inter-cohort (**D**) analyses net AIM^+^CD8^+^ T cell responses. **(EF)** Longitudinal (**E**) and inter-cohort (**F**) analyses of the net magnitude of cytokine^+^CD4^+^ T cell responses. The bold lines in A, C and E represent median values. The bars in B, D and F represent median ± interquartile ranges. In (**A, C, E**), the right panel show statistical comparisons using Wilcoxon tests. In **(B, D, F**), Mann-Whitney tests are shown. **(GH)** Heatmap showing temporal correlations of **(G)** AIM^+^CD4^+^ and **(H)** AIM^+^CD8^+^ T cells between the different time points for the two cohorts. The numbers in high square represent the correlation coefficient r. Significant Spearman tests results are indicated by stars (*: *p* < 0.1, **: *p* < 0.05, ***: *p* < 0.001). In **A-H**) LI cohort: n=26, SI cohort: n=12.

The AIM assay involved a 15-h incubation of PBMCs with an overlapping peptide pool spanning the Spike coding sequence of the ancestral strain and the measurement of CD69, CD40L, 4-1BB and OX40 upregulation upon stimulation. We used an AND/OR Boolean combination gating to assess total frequencies of antigen-specific CD4^+^ and CD8^+^ T cells (Figure S2AB) (Niessl et al., 2020a; Tauzin *et al*., 2021b). At D2, all individuals had detectable CD4^+^ T cell responses (Figure S2C), and most had measurable CD8^+^ T cell responses (Figure S2D).

AIM CD4^+^ T cell responses in the two cohorts differed at baseline and after the first dose (Figure 2A), suggesting a different pre-exposition to cross-reactive viruses participants of various age living in geographically distinct locations (Mateus et al., 2020). However, the responses converged after the second dose and no significant differences in AIM^+^CD4^+^ T cell responses were observed neither at D2 nor M2 (Figure 2B).

The trajectories of AIM^+^CD8^+^ T responses were heterogeneous. As reported in our previous study (Nayrac *et al*., 2022), LI participants elicited weak but significant responses after priming, a trend for stronger responses after the boost and contraction at M2 (Figure 3C). Consistent with AIM^+^CD4^+^ T cell responses, AIM^+^CD8^+^ T cell responses in the SI cohort was lower at baseline and D1 (Figure 3D). Nevertheless, we observed in this cohort a significant initial response priming, which did not significantly increase after the second dose. This plateau was further maintained at M2 at levels comparable to the post-attrition levels seen in the LI cohort, again indicating a convergence between the two vaccine modalities.

**Figure 3:**
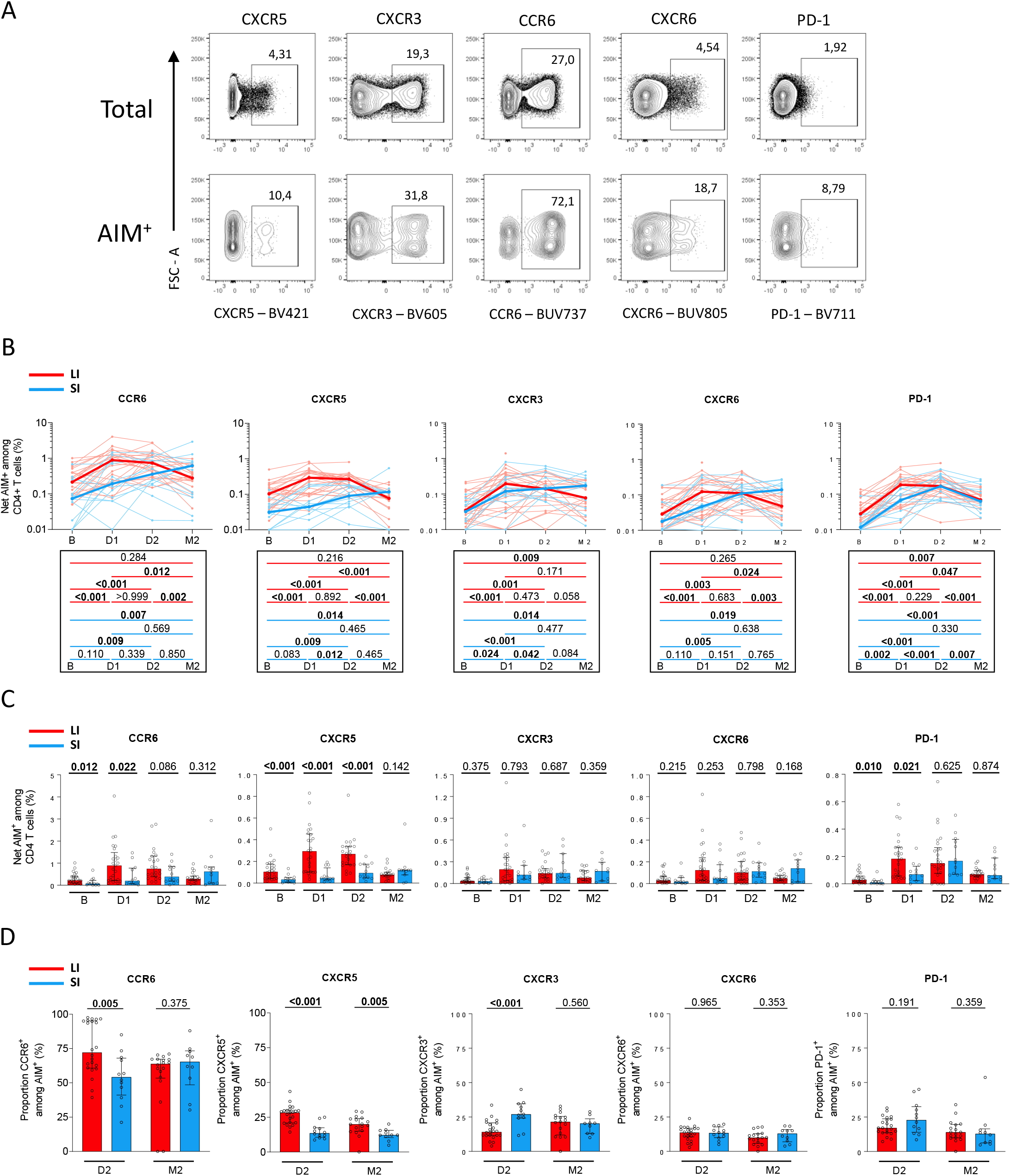
The second dose leads to convergence of some CD4^+^ T helper differentiation features differing early between the LI and SI cohorts. **(A)** Representative flow cytometry dot plots for the indicated univariate phenotypic population. **(B)** Net longitudinal frequencies of each AIM^+^CD4^+^ T cell subpopulation. Bold lines represent cohort’s median value. Bottom panel: Wilcoxon tests for each pairwise comparison. **(C)** Cohort comparisons at each time point for the subsets presented in (**A**). The bars represent median ± interquartile ranges. Mann-Whitney tests are shown. (**D**) Proportion of CXCR5^+^, CXCR3^+^, CCR6^+^, CXCR6^+^ and PD-1^+^ cells in total AIM^+^ CD4^+^ T cells at the D2 and M2 time points following the second dose. Mann-Whitney tests are shown. (**B-D**) LI cohort: n=26, SI cohort: n=12.

The ICS assay involved a 6-h stimulation with the Spike peptide pool and measurement of the effector molecules IFN-γ, IL-2, TNF-α, IL-17A, IL-10 and CD107a. We defined cytokine^+^CD4^+^ T cell responses by an AND/OR Boolean gating strategy (Figure S2E). Cytokine^+^ CD4^+^ T effector cells were readily detected after vaccination in most participants (Figure S2F). Total cytokine^+^CD8^+^ T cell responses were weak or undetectable in most participants, precluding their detailed analysis (Figure S2G). The ICS patterns in both cohorts paralleled the AIM assays, albeit at a lower magnitude (Figure 2EF). As for the AIM assay, cytokine responses converged at D2 and remained similar at M2. In contrast to AIM, however, cytokine^+^CD4^+^ T cell responses at M2 remained significantly higher than at baseline (Figure 2EF), showing longer-term memory poised for exerting effector functions.

As expansion of previously primed antigen-specific T cells may impact T cell responses to vaccination, we examined correlations across visits (Figure 2GH). We found in the LI cohort weak associations between post-priming AIM^+^ CD4^+^ T cell responses and those measured after boost or at the memory time point, respectively (Figure 2G). These associations were stronger in SI participants. We also found temporal associations for Spike-specific CD8^+^ T responses despite their lower magnitudes (Figure 2H).

These data show that in contrast to B cell responses, the initial differences in magnitude of Spike-specific CD4^+^ and CD8^+^ T cell responses that we observed between cohorts prior and early after priming disappeared after the second dose. The similar responses at the memory time point suggest that the time interval between the two doses has limited impact on the emergence and maintenance of Spike-specific CD4^+^ and CD8^+^ T cell immunity.

### The second dose leads to convergence of some CD4^+^ T helper differentiation features differing early between the LI and SI cohorts

As the interval had limited impact on the generation of CD4^+^ T cells but B cell responses remained lower after the second dose, we tested if different intervals could qualitatively affect CD4^+^ T cell responses and compared key CD4^+^ T cell subsets at D2 and M2 (Figure 3). We examined chemokine receptors that are preferentially, but not exclusively, expressed by some lineages and are involved in tissue homing (CXCR5 for Tfh; CXCR3 for Th1; CCR6 for Th17 and Th22 and mucosal homing; CXCR6 for pulmonary mucosal homing (Day et al., 2009; Morgan et al., 2015), and PD-1 as inhibitory checkpoint (Figure 3A), and assessed their longitudinal fluctuations (Figure 3BC).

CCR6^+^ cells were dominant in both cohorts, representing a median of 72% in LI and 54% in SI of all D2 responses, but with a wide inter individual variation (Figure 3D). CXCR5^+^, CXCR3^+^ and PD-1^+^ cells represented similar proportion of AIM^+^ CD4 T cells at D2. Median CXCR5^+^ were 28% (LI) and 14% (SI), median CXCR3^+^ were 14% (LI) and 27% (SI), and PD-1^+^ were 17% (LI) and 23% (SI). CXCR6^+^ cells were the rarest tested polarization, representing 13% (LI) and 14% (SI) of AIM^+^ CD4^+^ T cells. We observed a variable contribution of these Thelper subsets to the differences in total magnitude of CD4^+^ T cell responses between the LI and SI cohorts at baseline and after priming (Figure 2BC). The CCR6^+^ and CXCR5^+^ subset showed major differences with increased frequencies in LI at D2, but convergence at M2, whereas the kinetics of the CXCR3^+^ and CXCR6^+^ subsets showed no significant differences at any time points in the two cohorts (Figure 3BC). The PD-1^+^ subset differed initially but exhibited similar magnitude after the second dose (Figure 3BC). As shown by the relative fraction of each subset in the total AIM^+^ CD4^+^ T cell populations, some qualitative differences were still present shortly after the second dose but mostly waned at the memory timepoint (Figure 3D).

These results show that although the LI and SI cohorts presented qualitative differences at baseline and after the priming dose, repeat inoculation led to mostly converging features at the memory time point after the second dose, this despite the interval difference between doses in the two cohorts.

### The long and short vaccination regimens elicit largely similar patterns of CD4^+^ T cell effector functions

We next compared effector functions by ICS at D2 and M2, focusing on IFN-γ, TNF-α, IL-2, IL-10, and CD107a. IFN-γ^+^ and IL-2^+^ CD4^+^ T cells contracted at M2 in both cohorts, whereas TNFα remained constant (Figure 4AB). CD107a selectively contracted at M2 in the LI cohort. Consequently, CD107^+^ CD4^+^ T cells were more frequently detectable in the SI cohort, albeit at low levels. IL-10 followed a similar trend of decline, the difference in levels at M2 did not reach significance due to high variability among individual. After the second dose, we did not detect any statistically significant differences in the qualitative functional profile of CD4^+^ T cell responses elicited by the long and short interval vaccination schedules, as illustrated by the relative fraction of each cytokine in the total ICS response (Figure 4C).

**Figure 4:**
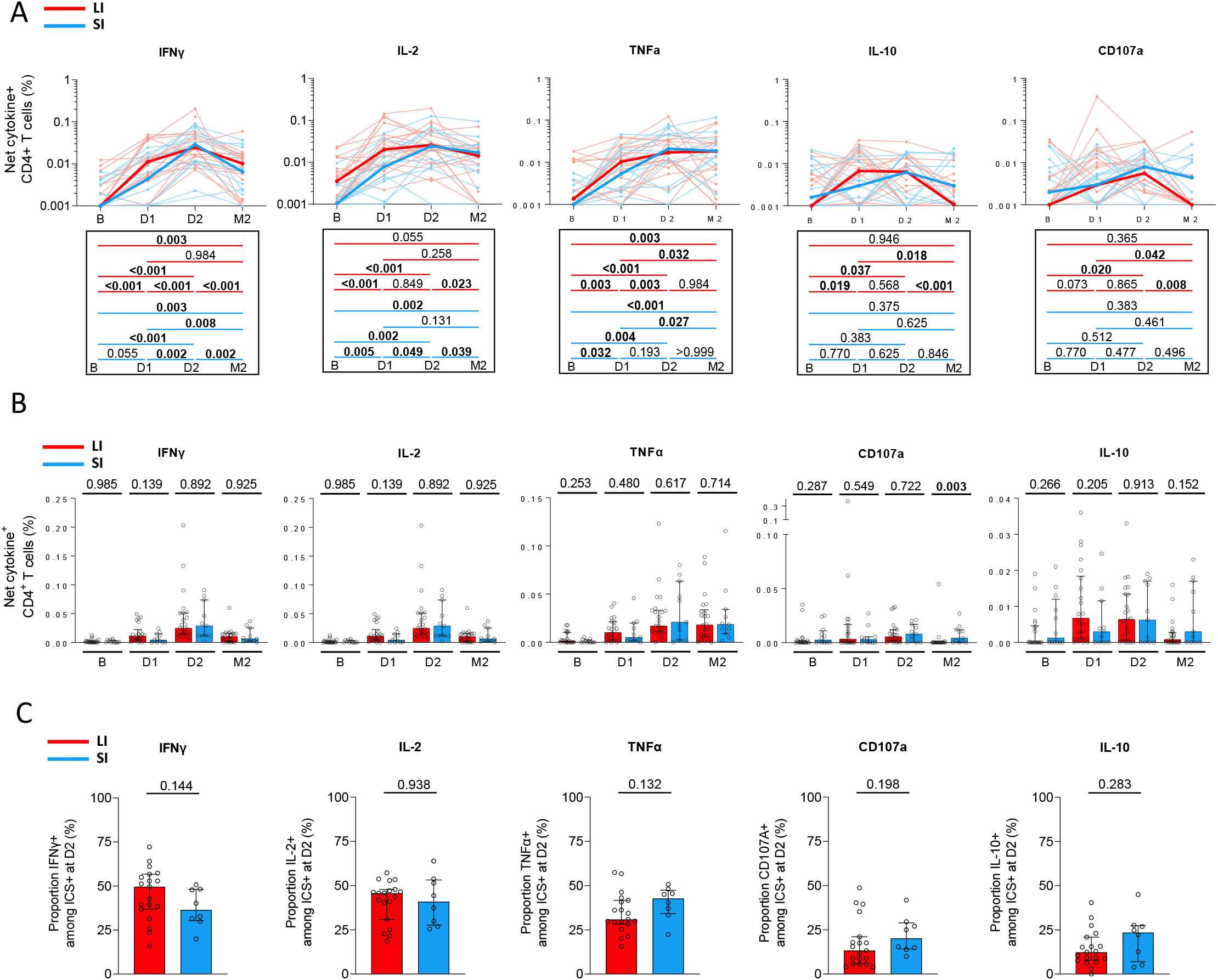
The long and short vaccination regimens elicit largely similar patterns of CD4^+^ T cell effector functions. **(A)** Longitudinal net frequencies of indicated cytokine^+^ CD4^+^ T cell subpopulations in the LI (red) and SI (blue) cohorts. Bold lines represent cohort’s median value. Lower panel: Wilcoxon tests for each pairwise comparison. **(B)** Cohort comparisons at each time point for each function represented in (**A**). The bars represent median ± interquartile ranges. Mann-Whitney tests are shown. (**C**) Proportions of IFN-γ, Il-2, TNF-α, IL-10 and CD107a-expressing cells among total cytokine^+^ CD4^+^ T cells. Mann-Whitney tests are shown to compare long and short-interval cohorts. (**AB**) LI cohort: n=26, SI cohort: n=12. (**C**) LI cohort: n=19, SI cohort: n=8.

Therefore, a longer interval between the first and second doses does not significantly alter the profile of tested effector CD4^+^ T functions.

## DISCUSSION

Several studies have shown that extending the interval between the first two doses of SARS-CoV-2 mRNA vaccines beyond the recommended regimens of 3 to 4 weeks can lead to stronger antibody responses (Grunau *et al*., 2021; Hall *et al*., 2022; Payne *et al*., 2021b; Tauzin *et al*., 2021a). These studies have led some public health agencies to modify their vaccination guidelines accordingly (e.g, 8 weeks or more between the primary two doses in Quebec (Quebec, 2022)). However, the impact of long-interval regimens on cellular immunity is still poorly known due to the paucity of studies performing side-by-side in-depth comparisons of different dosing regimens with the same assays. Here, we compared the antigen-specific B cell, CD4^+^ T cell and CD8^+^ T cell responses elicited in SARS-CoV-2 naïve participants by a 16-week interval regimen compared to the standard 3-4 weeks schedule. We observed that a long interval increased the magnitude and maturation of RBD-specific B cell responses, while completion of the primary vaccine series led to quantitatively and qualitatively similar memory CD4^+^ and CD8^+^ T cell memory responses in both regimens.

The RBD-specific B cells responses to the first vaccine dose were very consistent between the two cohorts, and did not appear impacted by the age difference between the groups. In contrast, the magnitude of these responses markedly differed after the second dose, with a robust increase in the LI cohort contrasting with a weak gain for the SI cohort. A second dose after a short interval might act like a prolonged antigen delivery rather than a recall of primed responses, thus explaining a more limited poor benefit. This difference persisted as a strong trend at the memory time point. These results are consistent with the fact that germinal centers remain active for several weeks after vaccination (Turner et al., 2021), with continuous evolution of the B cell compartment for several months (Cho et al., 2021) and accumulation of somatic hypermutations (Kim et al., 2022; Paschold et al., 2022; Turner et al., 2021). Hence, an early second dose likely corresponds to a suboptimal timing in terms of re-exposure to the cognate antigen, while a longer interval allows for a better evolution of the B cell repertoire. In line with these findings, the B cell maturation profile differed between the LI and SI cohorts after the second dose: almost all RBD-specific B cells presented an isotype-switched IgG^+^ phenotype in LI participants, contrasting with a sizeable minority of IgM^+^ cells in SI volunteers. The memory differentiation phenotype was also consistent with this profile, with a larger fraction of RBD^+^ B cells with a CD27^+^ IgD-memory phenotype early after boost in the LI participants. As we previously reported (Nayrac *et al*., 2022) the RBD-specific B cell responses were dominated by the double-negative CD27^-^ IgD^-^ cells, including at the memory time point. This phenotypic subset was described in autoimmune diseases (Jenks et al., 2018; Wei et al., 2007) and in response to vaccination (Ruschil et al., 2020). We observed some cells with a naïve-like phenotype based on these two markers (CD27^-^ IgD^+^) after the second dose in the SI cohort, but almost none in the LI cohort. These cells were also absent at baseline and in previously-infected individuals (Nayrac et al., 2022), suggesting recently activated B cells. Taken together, these results suggest that the long interval regimen is beneficial to the generation and maturation of the B cell compartment.

We observed that Spike-specific CD4^+^ and CD8^+^ T cell responses at baseline were significantly stronger in the LI compared to the SI cohort. Other studies have shown that cross-reactive immunity to other coronaviruses play a major role in shaping these pre-existing SARS-CoV-specific CD4^+^ and CD8^+^ T cell responses (Braun et al., 2020; Grifoni et al., 2020; Mateus *et al*., 2020; Shrock et al., 2020). Of note, the two cohorts were of geographically distinct locations (LI: Montreal, SI: Philadelphia) and the LI participants were significantly older than the SI volunteers. While the lack of sufficient PBMC samples precluded direct testing of cross-reactivity, it is therefore likely that differential previous exposure to endemic coronaviruses contributes to the pre-vaccination differences observed. These differences persisted after the first vaccine dose, consistent with previously reported association between pre-existing T cell immunity and responses after priming (Grifoni et al., 2020; Le Bert et al., 2020; Loyal et al., 2021; Tauzin *et al*., 2021b). Importantly, however, the quantitative and qualitative differences in CD4^+^ and CD8^+^ T cell responses decreased already early after the second vaccine dose and waned almost entirely at the memory time point collected 10 to 16 weeks after the boost. This convergence was present both in phenotypic AIM assays (e.g, for CXCR5^+^ and CCR6^+^ CD4^+^ T cells) and functional ICS assays. IFN-γ^+^ and IL-2^+^ CD4^+^ T cell responses were comparable in the two cohorts, consistent with a recent study (Hall et al., 2022). Similarly, we did not identify differences in memory responses for TNF-α and IL-10 production. The small difference in CD107a^+^ CD4^+^ T cells frequencies should be interpreted with caution, given the very low magnitude of these responses. At first sight, our IL-2 data differ from another study that reported stronger memory IL-2^+^ CD4 T cell responses in long-interval vaccination (Payne *et al*., 2021a). However, the timeline may contribute to these differences. In our study, we assessed memory later after the second dose (10-16 weeks versus 4 weeks in (Payne et al., 2021a)). Therefore, completion of the primary 2-dose vaccination leads to convergent CD4^+^ and CD8^+^ T cell memory responses irrespective of dosing interval.

While the initial rationale of delaying the second dose was to provide some level of immunity more rapidly to a larger number of people in the context of limiting vaccine supply, our results show that this strategy is beneficial to the generation of B cell responses without negative impact on T cell immunity. The potential immunological benefits of increasing the interval between doses must be weighed against a prolonged window of suboptimal protection, particularly while the virus and its different variants of concern are circulating in the population. Many countries now recommend a third dose, and more, although compliance with additional inoculations is a significant issue. Whether additional inoculations further abrogate the differences in cellular immunity observed between the long and short interval regimens after the primary vaccination series warrants further investigation.

## Limitations of the study

Here, we focused the investigations on cellular responses. We previously investigated the impact of the dosing interval on humoral responses and showed a benefit of a longer interval on the quality of antibody immunity (Chatterjee et al., 2022).

Here, we investigated individuals who were SARS-CoV-2 naïve prior to vaccination. However, we did not investigate the impact of a long versus short-interval vaccine regimens in previously-infected people. Further comparative studies are therefore required to assess the impact of dosing interval on cellular hybrid immunity.

The demographically distinct LI and SI cohorts presented differences in T cell responses at baseline that we interpreted this as likely reflecting the presence of a pre-existing pool of cross-reactive cells to other coronaviruses. Formal demonstration would require epitope-specific mapping of T cell responses, for which we did not have enough PBMC samples available.

We analyzed the cellular responses to ancestral strain antigens corresponding to the mRNA vaccines. The limiting availability in PBMC did not allow to assess the impact of dosing interval on B and T cell responses to variants of concern.

The size of the cohorts investigated here, particularly of the short-interval group, is not sufficient to uncover potential smaller qualitative differences in the T cell responses that might be cause by different intervals. However, the contrasting results obtained for B cell responses compared to T cell responses are clear enough to conclude that modifying the time between vaccine inoculations has a much bigger impact on B cell than T cell immunity.

Our study conducted in a low-risk HCW cohort may not be generalizable to vulnerable groups, particularly immunocompromised or elderly populations, in which the cellular immune responses and the risk/benefit ratio may differ. Future studies will be required to better quantify the immune response over time in these populations.

## ACKNOWLEDGMENTS

The authors are grateful to the study participants. We thank the CRCHUM BSL3 and Flow Cytometry Platforms for technical assistance, Dr. Johanne Poudrier for advice and discussions. This work was supported by a FRQS Merit Research Scholar award to D.E.K, the Fondation du CHUM, le Ministère de l’Économie et de l’Innovation du Québec, Programme de soutien aux organismes de recherche et d’innovation (to A.F), a CIHR operating grant # 178344 (D.E.K and A.F), a foundation grant #352417 (A.F), a CIHR operating Pandemic and Health Emergencies Research grant #177958 (A.F), and an Exceptional Fund COVID-19 from the Canada Foundation for Innovation (CFI) #41027 to A.F and D.E.K. The Symphony flow cytometer was funded by a John R. Evans Leaders Fund Leader Fund from the Canada Foundation for Innovation (# 37521 to D.E.K) and the Fondation Sclérodermie Québec. A.F. is the recipient of Canada Research Chair on Retroviral Entry no. RCHS0235 950-232424. V.M.L. is supported by a FRQS Junior 1 salary award, G.S by scholarship from the Department of Microbiology, Infectious Disease and Immunology of the University of Montreal. This work was also supported by NIH funds: grants AI108545, AI155577, AI149680, and U19AI082630 (to E.J.W.), the University of Pennsylvania Perelman School of Medicine COVID Fund (to R.R.G. and E.J.W.); the University of Pennsylvania Perelman School of Medicine 21st Century Scholar Fund (to R.R.G.); and the Paul and Daisy Soros Fellowship for New Americans (to R.R.G). The funders had no role in study design, data collection and analysis, decision to publish, or preparation of the manuscript.

## AUTHOR CONTRIBUTIONS

A.N., G.S. M.D, A.F. and D.E.K. designed the studies. A.N., G.S, M.N, N.B and M.L. performed B cell and T cell assays. A.N, G.S. and M.D. performed and analyzed the B and T cell experiments. J.N. contributed to the T cell assay design. H.M., L.G., C.M., P.A., C.T. J.C.W. and V.M.L. secured and processed blood samples. M.M.P., R.R.G., A.R.G, E.J.W. provided unique reagents. G.G. produced and purified proteins. M.D., A.N, G.S. and D.E.K. wrote the manuscript. Every author has read, edited and approved the final manuscript.

## DECLARATION OF INTERESTS

A.R.G. is a consultant for Relation Therapeutics. E.J.W. is consulting for or is an advisor for Merck, Marengo, Janssen, Related Sciences, Synthekine, and Surface Oncology. E.J.W. is a founder of Surface Oncology, Danger Bio, and Arsenal Biosciences. The other authors have no conflict of interest to declare.

## STAR METHODS

### RESOURCE AVAILABILITY

#### Lead contact

Further information and requests for resources and reagents should be directed to and will be fulfilled by the lead contact, Daniel E. Kaufmann.

#### Materials availability

All unique reagents generated during this study are available from the Lead contact upon a material transfer agreement (MTA).

#### Data and code availability

The published article includes all datasets generated and analyzed for this study. Further information and requests for resources and reagents should be directed to and will be fulfilled by the Lead Contact Author.

### EXPERIMENTAL MODEL AND SUBJECT DETAILS

#### Ethics Statement

All work was conducted in accordance with the Declaration of Helsinki in terms of informed consent and approval by an appropriate institutional board. Blood samples were obtained from donors who consented to participate in this research project at CHUM (19.381). Individuals from the Philadelphia were enrolled in the study with approval from the University of Pennsylvania Institutional Review Board (IRB# 844642). All participants were otherwise healthy and did not report any history of chronic health conditions.

#### Participants

No specific criteria such as number of patients (sample size), clinical or demographic were used for inclusion, beyond negative PCR confirmation for SARS-CoV-2 infection. The study was conducted in 26 SARS-CoV-2 naïve individuals with a long interval, and 12 with a short interval. All the information is summarized in Table 1.

#### PBMCs collection

PBMCs were isolated from blood samples by Ficoll density gradient centrifugation and cryopreserved in liquid nitrogen until use.

### METHOD DETAILS

#### Protein expression and purification

FreeStyle 293F cells (Thermo Fisher Scientific) were grown in FreeStyle 293F medium (Thermo Fisher Scientific) to a density of 1 × 10^6^ cells/mL at 37°C with 8 % CO_2_ with regular agitation (150 rpm). Cells were transfected with a plasmid coding for SARS-CoV-2 S RBD using ExpiFectamine 293 transfection reagent, as directed by the manufacturer (Invitrogen) (Beaudoin-Bussieres et al., 2020; Prevost et al., 2020). One week later, cells were pelleted and discarded. Supernatants were filtered using a 0.22 µm filter (Thermo Fisher Scientific). The recombinant RBD proteins were purified by nickel affinity columns, as directed by the manufacturer (Thermo Fisher Scientific). The RBD preparations were dialyzed against phosphate-buffered saline (PBS) and stored in aliquots at -80°C until further use. To assess purity, recombinant proteins were loaded on SDS-PAGE gels and stained with Coomassie Blue.

#### SARS-CoV-2-specific B cells characterization

To detect SARS-CoV-2-specific B cells, we conjugated recombinant RBD proteins with Alexa Fluor 488 or Alexa Fluor 594 (Thermo Fisher Scientific) according to the manufacturer’s protocol. 2 × 10^6^ frozen PBMC from SARS-CoV-2 naïve and previously-infected donors were prepared in Falcon® 5ml-round bottom polystyrene tubes at a final concentration of 4 × 10^6^ cells/mL in RPMI 1640 medium (GIBCO) supplemented with 10% of fetal bovine serum (Seradigm), Penicillin-Streptomycin (GIBCO) and HEPES (GIBCO). After a rest of 2h at 37°C and 5% CO_2_, cells were stained using Aquavivid viability marker (GIBCO) in DPBS (GIBCO) at 4°C for 20min. The detection of SARS-CoV-2-antigen specific B cells was done by adding the RBD probes to the antibody cocktail listed in Table S1. Staining was performed at 4°C for 30min and cells were fixed using 2% paraformaldehyde at 4°C for 15min. Stained PBMC samples were acquired on Symphony cytometer (BD Biosciences) and analyzed using FlowJo v10.8.0 software.

#### Activation-induced marker (AIM) assay

The AIM assay (Morou et al., 2019; Niessl *et al*., 2020a; Niessl et al., 2020b; Tauzin *et al*., 2021b) was adapted for SARS-CoV-2 specific CD4 and CD8 T cells was as previously described (Tauzin *et al*., 2021b). PBMCs were thawed and rested for 3h in 96-well flat-bottom plates in RPMI 1640 supplemented with HEPES, penicillin and streptomycin and 10% FBS. 1.7×10^6^ PBMCs were stimulated with a S glycoprotein peptide pool (0.5 μg/ml per peptide, corresponding to the pool of 315 overlapping peptides (15-mers) spanning the complete amino acid sequence of the Spike glycoprotein (JPT) for 15h at 37 °C and 5% CO_2_. CXCR3, CCR6, CXCR6 and CXCR5 antibodies were added in culture 15 min before stimulation. A DMSO-treated condition served as a negative control and *Staphylococcus enterotoxin B* SEB-treated condition (0.5 μg/ml) as positive control. Cells were stained for viability dye for 20 min at 4 °C then surface markers (30 min, 4 °C). Abs used are listed in the Table S2. Cells were fixed using 2% paraformaldehyde for 15min at 4 °C before acquisition on Symphony cytometer (BD Biosciences). Analyses were performed using FlowJo v10.8.0 software.

#### Intracellular Cytokine Staining (ICS)

The ICS assay adapted to study SARS-CoV-2-specific T cells was previously described (Tauzin *et al*., 2021b). PBMCs were thawed and rested for 2 h in RPMI 1640 medium supplemented with 10% FBS, Penicillin-Streptomycin (Thermo Fisher scientific, Waltham, MA) and HEPES (Thermo Fisher scientific, Waltham, MA). 1.7×10^6^ PBMCs were stimulated with a S glycoprotein peptide pool (0.5 μg/mL per peptide from JPT, Berlin, Germany) corresponding to the pool of 315 overlapping peptides (15-mers) spanning the complete amino acid sequence of the S glycoprotein.

Cell stimulation was carried out for 6h in the presence of mouse anti-human CD107a, Brefeldin A and monensin (BD Biosciences, San Jose, CA) at 37 °C and 5% CO_2_. DMSO-treated cells served as a negative control, and SEB as positive control. Cells were stained for Aquavivid viability marker (Thermo Fisher scientific, Waltham, MA) for 20 min at 4 °C and surface markers (30 min, 4 °C), followed by intracellular detection of cytokines using the IC Fixation/Permeabilization kit (Thermo Fisher scientific, Waltham, MA) according to the manufacturer’s protocol before acquisition on a Symphony flow cytometer (BD Biosciences) and analysis using FlowJo v10.8.0 software. Abs used are listed in the Table S3.

Characterization of effector functions among total cytokine^+^ cells, defined by our ORgate strategy, was conducted on donors with >5 cytokine^+^ cells that represented a two-fold increase over the unstimulated condition to avoid biaised phenotyping. Given these criteria, only D2 could be analyzed.

### QUANTIFICATION AND STATISTICAL ANALYSIS

#### Statistical analysis

Symbols represent biologically independent samples of HCW from LI and SI cohorts. Lines connect data from the same donor. Thick lines represent median values. Differences in responses for the same patient before and after vaccination were performed using Wilcoxon matched pair tests. Differences in responses between individuals from LI and SI cohorts were measured by Mann-Whitney tests. Wilcoxon and Mann-Whitney tests were generated using GraphPad Prism version 8.4.3 (GraphPad, San Diego, CA) (Rodda *et al*., 2022).

P values < 0.05 were considered significant. P values are indicated for each comparison assessed. For descriptive correlations, Spearman’s R correlation coefficient was applied. For graphical representation on a log scale (but not for statistical tests), null values were arbitrarily set at the minimal values for each assay.

#### Software scripts and visualization

Graphics and pie charts were generated using GraphPad PRISM version 8.4.1 and ggplot2 (v3.3.3) in R (v4.1.0).

## SUPPLEMENTAL INFORMATION

Table S1. Flow cytometry antibody staining panel for B cells characterization, related to the STAR Methods section.

Table S2. Flow cytometry antibody staining panel for activation-induced marker assay, related to the STAR Methods section.

Table S3. Flow cytometry antibody staining panel for intracellular detection, related to the STAR Methods section.

