## Supplemental Figures and Tables for "A long interval between priming and boosting SARS-CoV-2 mRNA vaccine doses enhances B cell responses with limited impact on T cell immunity"

### Supplemental 1

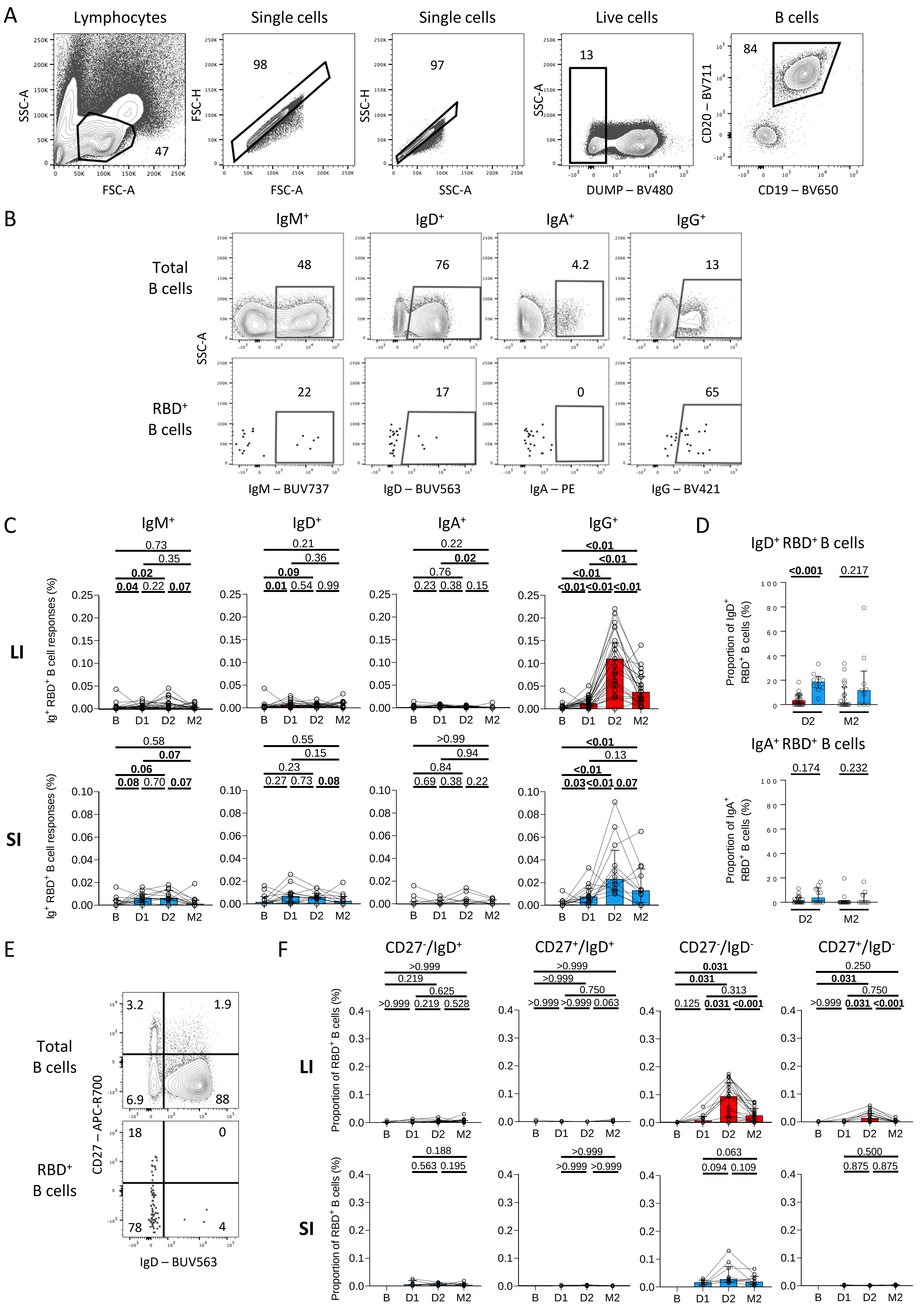

Supplemental 2

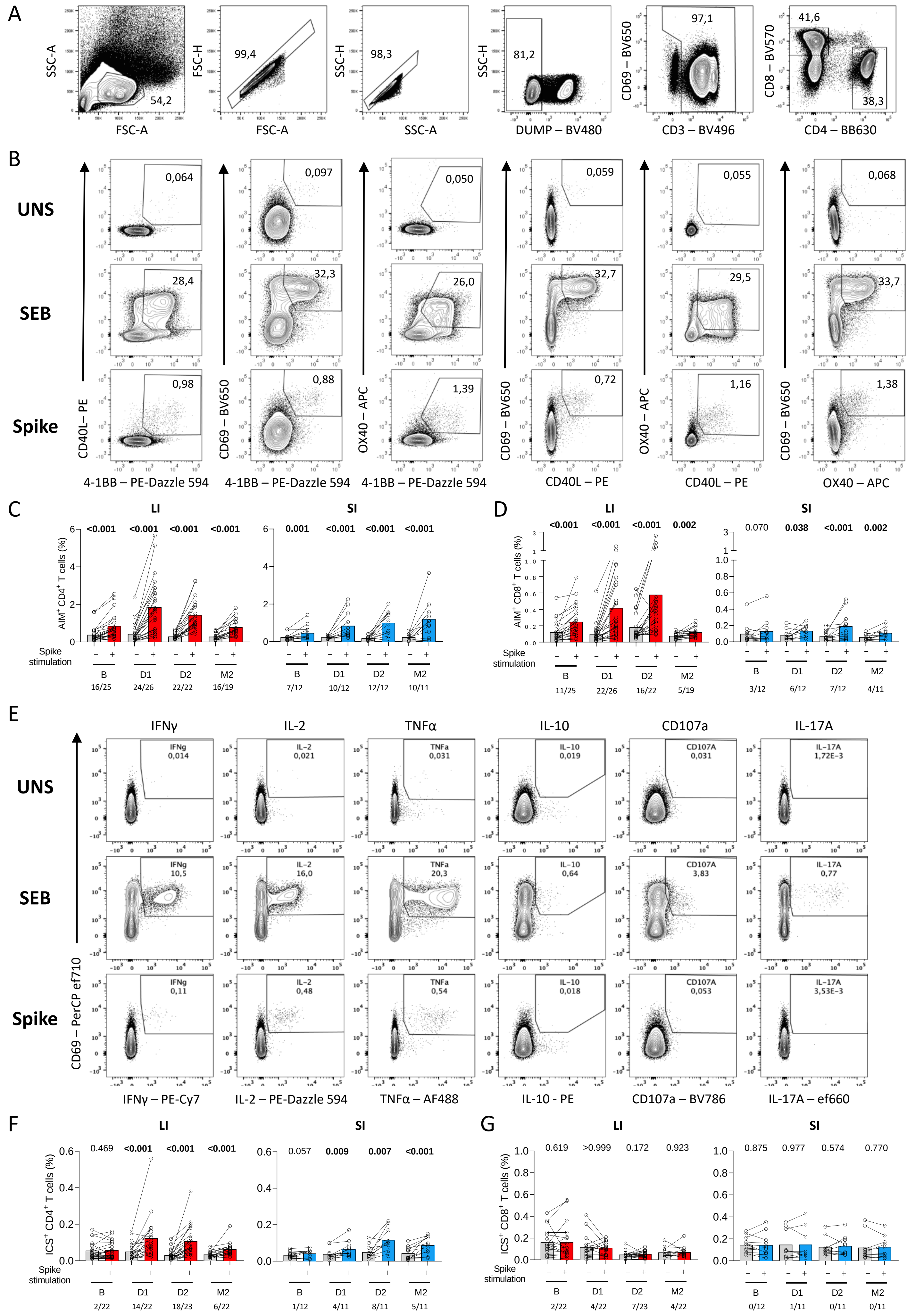

#### SUPPLEMENTARY FIGURE LEGENDS

**Figure S1. Related to figure 1. (A)** Gating strategy to identify RBD-specific B cell responses. (**B**) Examples of gatings for IgM, IgD, IgA and IgG expression on total $CD19^{+}CD20^{+}$ B cells or RBD-specific B cells. **(C)** Longitudinal trajectories of isotype expression frequencies in long interval (LI) (n=26) and short interval (SI) (n=12) participants. Lines connect data points for individual participants. Wilcoxon tests are shown above each panel. **(D)** Proportion of $IgD^{+}$ $RBD^{+}$ B cells and $IgA^{+}$ $RBD^{+}$ B cells at D2 and M2 in LI (red, n=26) and SI (blue, n=12) participants. Mann-Whitney tests are shown. **(E)** Example of the gating strategy of IgD and CD27 co-expression on RBD-specific B cells. **(F)** Longitudinal frequency of each IgD and CD27 RBD-B phenotypes in $CD19^{+}$ $CD20^{+}$ B cells for LI (n=26) and SI (n=12) participants. Wilcoxon tests are shown above.

**Figure S2: Related to figure 2. (A)** Representative upstream generic gating and **(B)** ORgate strategy to identify SARS-CoV-2-specific $AIM^{+}$ T cells. For simplicity, the example focuses on $CD4^{+}$ T cells. **(CD)** Raw frequencies of **(C)** $AIM^{+}$ $CD4^{+}$ and **(D)** $CD8^{+}$ T cells following *ex vivo* stimulation of PBMCs with a pool of SARS-CoV-2 Spike peptides. As a control, PBMCs cells were left unstimulated (grey bars). The data for LI (red, n=26) and SI (blue, n=12) individuals are displayed. The bars represent median values. Wilcoxon tests are shown. The number of conditions reaching >2x no Ag are shown below each timepoint. **(E)** Representative ORgate strategy to identify SARS-CoV-2-specific cytokine-expressing T cells. For simplicity, the example focuses on $CD4^{+}$ T cells. **(FG)** Raw frequencies of cytokine-expressing (**F**) $CD4^{+}$ and (**G**) $CD8^{+}$ T cells following *ex vivo* stimulation of PBMCs with a pool of SARS-CoV-2 Spike peptides. As a control, PBMCs cells were left unstimulated (grey bars). The data for LI (red, n=26) and SI (PI; blue, n=12) individuals are displayed. The bars represent median values. Wilcoxon tests are shown. The number of conditions reaching >2x no Ag are shown below each time point.

**Table S1. Flow cytometry antibody staining panel for B cells characterization**

| Marker-Fluorophore | Clone | Source | Catalog # |
|---|---|---|---|
| CD3 – BV480 | UCHT1 | BD Biosciences | 566105 |
| CD14 – BV480 | M5E2 | BD Biosciences | 746304 |
| CD16 – BV480 | 3G8 | BD Biosciences | 566108 |
| CD19 – BV650 | SJ25C1 | Biolegend | 363026 |
| CD20 – BV711 | 2H7 | Biolegend | 563126 |
| CD21 – BV786 | B-LY4 | BD Biosciences | 740969 |
| CD24 – BUV805 | ML5 | BD Biosciences | 742010 |
| CD27 – APC-R700 | M-T271 | BD Biosciences | 565116 |
| CD38 – BB790 | HIT2 | BD Biosciences | CUSTOM |
| CD56 – BV480 | NCAM16.2 | BD Biosciences | 566124 |
| CD138 – BUV661 | MI15 | BD Biosciences | 5 749873 |
| CCR10 – BUV395 | 1B5 | BD Biosciences | 565322 |
| HLA-DR – BB700 | G46-6 | BD Biosciences | 566480 |
| IgA - PE | IS11-8E10 | Miltenyi | 130-113-476 |
| IgD – BUV563 | IA6-2 | BD Biosciences | 741394 |
| IgG – BV421 | G18-147 | BD Biosciences | 562581 |
| IgM – BUV737 | UCH-B1 | Thermo Fisher Scientific | 748928 |
| LIVE/DEAD Fixable dead cell | N/A | Thermo Fisher Scientific | L34960 |

**Table S2. Flow cytometry antibody staining panel for activation-induced marker assay**

| Marker-Fluorophore | Clone | Source | Catalog # |
|---|---|---|---|
| CD3 – BUV496 | UCHT1 | BD | 612941 |
| CD4 – BB630 | SK3 | BD | 624294 |
| CD8 – BV570 | RPA-T8 | Biolegend | 301037 |
| CD14 – BV480 | M5E2 | BD | 746304 |
| CD19 – BV480 | HIB19 | BD | 746457 |
| CD38 – BB790 | HIT2 | BD | CUSTOM |
| CD45RA – PerCP Cy5.5 | HI100 | BD | 563429 |
| CD69 – BV650 | FN50 | Biolegend | 310934 |
| CD134 (OX40) - APC | ACT35 | BD | 563473 |
| CD137 (4-1BB) – PE-Dazzle 594 | 4B4-1 | Biolegend | 309826 |
| CD154 (CD40L) - PE | TRAP1 | BD | 555700 |
| CD183 (CXCR3) – BV605 | G025H7 | Biolegend | 353728 |
| CD185 (CXCR5) – BV421 | J25D4 | Biolegend | 356920 |
| CD186 (CXCR6) – BUV805 | 13B 1E5 | BD | 748448 |
| CD196 (CCR6) – BUV737 | 11A9 | BD | 564377 |
| CD279 (PD1) – BV711 | EH122H | Biolegend | 329928 |
| HLA-DR - FITC | LN3 | Biolegend | 327005 |
| LIVE/DEAD Fixable dead cell | N/A | Thermo Fisher Scientific | L34960 |

**Table S3. Flow cytometry antibody staining panel for intracellular cytokines staining assay**

| Marker-Fluorophore | Clone | Source | Catalog # |
|---|---|---|---|
| CD3 – BUV395 | UCHT1 | BD Biosciences | 563546 |
| CD4 – BV711 | L200 | BD Biosciences | 563913 |
| CD8 – BV570 | RPA-T8 | Biolegend | 301037 |
| CD14 – BUV805 | M5E2 | BD Biosciences | 612902 |
| CD16 – BV650 | 3G8 | Biolegend | 302042 |
| CD19 – APC-eFluor780 | HIB19 | Thermo Fisher Scientific | 47-0199 |
| CD56 – BUV737 | NCAM16.2 | BD Biosciences | 564448 |
| CD69 – PerCP-eFluor710 | FN50 | Thermo Fisher Scientific | 46-0699-42 |
| CD107A – BV786 | H4A3 | BD Biosciences | 563869 |
| IFN-γ – PECy7 | B27 | BD Biosciences | 557643 |
| CD154 (CD40L) – BV421 | TRQP1 | BD Biosciences | 563886 |
| IL-2 – PE-Dazzle 594 | MQ1-17H12 | Biolegend | 500344 |
| IL-10 - PE | JES3-9D7 | BD Biosciences | 554498 |
| IL-17A – eFluor660 | eBio64CAP17 | Thermo Fisher Scientific | 50-7179-42 |
| TNF-α – Alexa Fluor 488 | Mab11 | Thermo Fisher Scientific | 502915 |
| Granzym B – Alexa Fluor 700 | GB11 | BD | 561016 |
| LIVE/DEAD Fixable dead cell | N/A | Thermo Fisher Scientific | L34960 |
